# Clusters of bacterial RNA polymerase are biomolecular condensates that assemble through liquid-liquid phase separation

**DOI:** 10.1101/2020.03.16.994491

**Authors:** A-M Ladouceur, B Parmar, S Biedzinski, J Wall, SG Tope, D Cohn, A Kim, N Soubry, R Reyes-Lamothe, SC Weber

## Abstract

Once described as mere “bags of enzymes”, bacterial cells are in fact highly organized, with many macromolecules exhibiting non-uniform localization patterns. Yet the physical and biochemical mechanisms that govern this spatial heterogeneity remain largely unknown. Here, we identify liquid-liquid phase separation (LLPS) as a mechanism for organizing clusters of RNA polymerase (RNAP) in *E. coli*. Using fluorescence imaging, we show that RNAP quickly transitions from a dispersed to clustered localization pattern as cells enter log phase in nutrient-rich media. RNAP clusters are sensitive to hexanediol, a chemical that dissolves liquid-like compartments in eukaryotic cells. In addition, we find that the transcription antitermination factor NusA forms droplets *in vitro* and *in vivo*, suggesting that it may nucleate RNAP clusters. Finally, we use single-molecule tracking to characterize the dynamics of cluster components. Our results indicate that RNAP and NusA molecules move inside clusters, with mobilities faster than a DNA locus but slower than bulk diffusion through the nucleoid. We conclude that RNAP clusters are biomolecular condensates that assemble through LLPS. This work provides direct evidence for LLPS in bacteria and suggests that this process serves as a universal mechanism for intracellular organization across the tree of life.

**Significance:** Bacterial cells are small and were long thought to have little to no internal structure. However, advances in microscopy have revealed that bacteria do indeed contain subcellular compartments. But how these compartments form has remained a mystery. Recent progress in larger, more complex eukaryotic cells has identified a novel mechanism for intracellular organization known as liquid-liquid phase separation. This process causes certain types of molecules to concentrate within distinct compartments inside the cell. Here, we demonstrate that the same process also occurs in bacteria. This work, together with a growing body of literature, suggests that liquid-liquid phase separation is a universal mechanism for intracellular organization that extends across the tree of life.

## Introduction

Living cells are divided into functional compartments called organelles. In eukaryotes, lipid membranes create a diffusion barrier between organelles and the cytoplasm, such that each compartment maintains a distinct biochemical composition that is tailored to its function (1). Prokaryotes typically lack internal membranes and instead must use alternate mechanisms for spatial and functional organization. For example, bacterial microcompartments, such as the carboxysome, are surrounded by a selectively-permeable protein shell (2). Storage granules also contain a surface layer of proteins (3). In addition to these discrete organelle-like structures, chromosomal loci (4-6), proteins (7, 8) and lipids (9) have all been found to exhibit non-uniform localization patterns in bacteria. However, in most cases, the mechanisms that give rise to such spatial heterogeneity, and its functional significance, remain unclear.

Liquid-liquid phase separation (LLPS) has recently emerged as a novel mechanism for compartmentalization in eukaryotic cells (10). This process mediates the assembly of “biomolecular condensates”, an unusual class of organelles that lack delimiting membranes (11). For example, P granules (12) and stress granules (13) consist of local concentrations of protein and RNA which rapidly assemble in the cytoplasm in response to developmental or environmental changes. The nucleoplasm also undergoes phase separation to generate a variety of “nuclear bodies” (14, 15), including the prominent nucleolus (16, 17). Intriguingly, membraneless bodies have also been described in chloroplasts (18, 19) and mitochondria (20). Given these organelles’ endosymbiotic origin (21), these observations raise the possibility that prokaryotes also use LLPS to compartmentalize their cells.

In eukaryotes, LLPS is mediated by proteins containing multivalent domains and/or disordered regions (22-24) that bring molecules together into dynamic condensates through transient interactions. Sequence analysis predicts that bacterial proteomes have a much lower frequency of disordered regions than eukaryotic proteomes: only 4.2% compared to 33%, respectively (25). This paucity of disorder raises doubts about the prevalence of LLPS in this domain of life. Nevertheless, recent work suggests that bacteria may indeed contain biomolecular condensates (26). For example, the DEAD-box helicases DeaD, SrmB and RhlE form foci in *E. coli* (27) and RNase E forms foci in *Caulobacter crescentus* and other α-proteobacteria (28). Moreover, the disordered C-terminal domain of RNase E is necessary and sufficient for foci assembly *in vivo*.

Despite these observations, direct evidence for LLPS in bacteria is still lacking. This gap arises in part due to the small size of bacteria and the inherent difficulty of analyzing structures near or below the diffraction limit. Yet a more fundamental problem also exists: a lack of consensus criteria to define LLPS *in vivo*. This is true even in eukaryotic systems, as highlighted by several recent reviews (15, 29, 30).

To overcome these challenges, we combine traditional chemical and genetic perturbations with single-molecule tracking to investigate the clustering of bacterial RNA polymerase (RNAP). RNAP is distributed throughout the nucleoid in cells grown in minimal media, but concentrates into distinct clusters in cells grown in rich media (31, 32). This nutrient-dependent localization has been well-characterized (33, 34), but the mechanism(s) that govern RNAP clustering remains unclear (35, 36). Here, we show that clusters of bacterial RNAP are biomolecular condensates. They assemble in a rapid and discontinuous fashion; they are sensitive to hexanediol; and their molecular components are dynamic. Our results demonstrate that LLPS is a universal organizing principle that extends across the tree of life.

## Results

### RNAP clusters during log phase in rich media

To investigate the mechanism(s) governing the spatial organization of RNAP in bacterial cells, we generated an *E. coli* strain that expresses an mCherry fusion of the β’ subunit (RpoC) from its endogenous locus. We performed outgrowth experiments by diluting saturated overnight cultures into fresh media at 37°C. RNAP clusters rapidly disperse when cells are transferred from liquid culture to agarose pads at room temperature (32, 37). To preserve the native subcellular distribution of RNAP, we collected samples at regular time intervals and immediately fixed them in formaldehyde before imaging. Fluorescence localization patterns observed in fixed cells were similar to those in live cells when imaged at 37°C (Fig. S1).

The spatial organization of RNAP depended on the phase of growth. RNAP is dispersed throughout the nucleoid in lag-phase cells (Fig. 1A, 0 min). In nutrient-rich media (LB and EZ), RNAP clusters into several bright foci per cell during log phase (Fig. 1A, 60-180 min) and gradually disperses as cells re-enter stationary phase (Fig. 1A, 240-360 min). In minimal media (M9), RNAP clustering is delayed and less pronounced (Fig. 1A, 120-300 min). These observations are consistent with previous reports examining log-phase cells grown in rich and minimal media (32-34).

**Figure 1.**
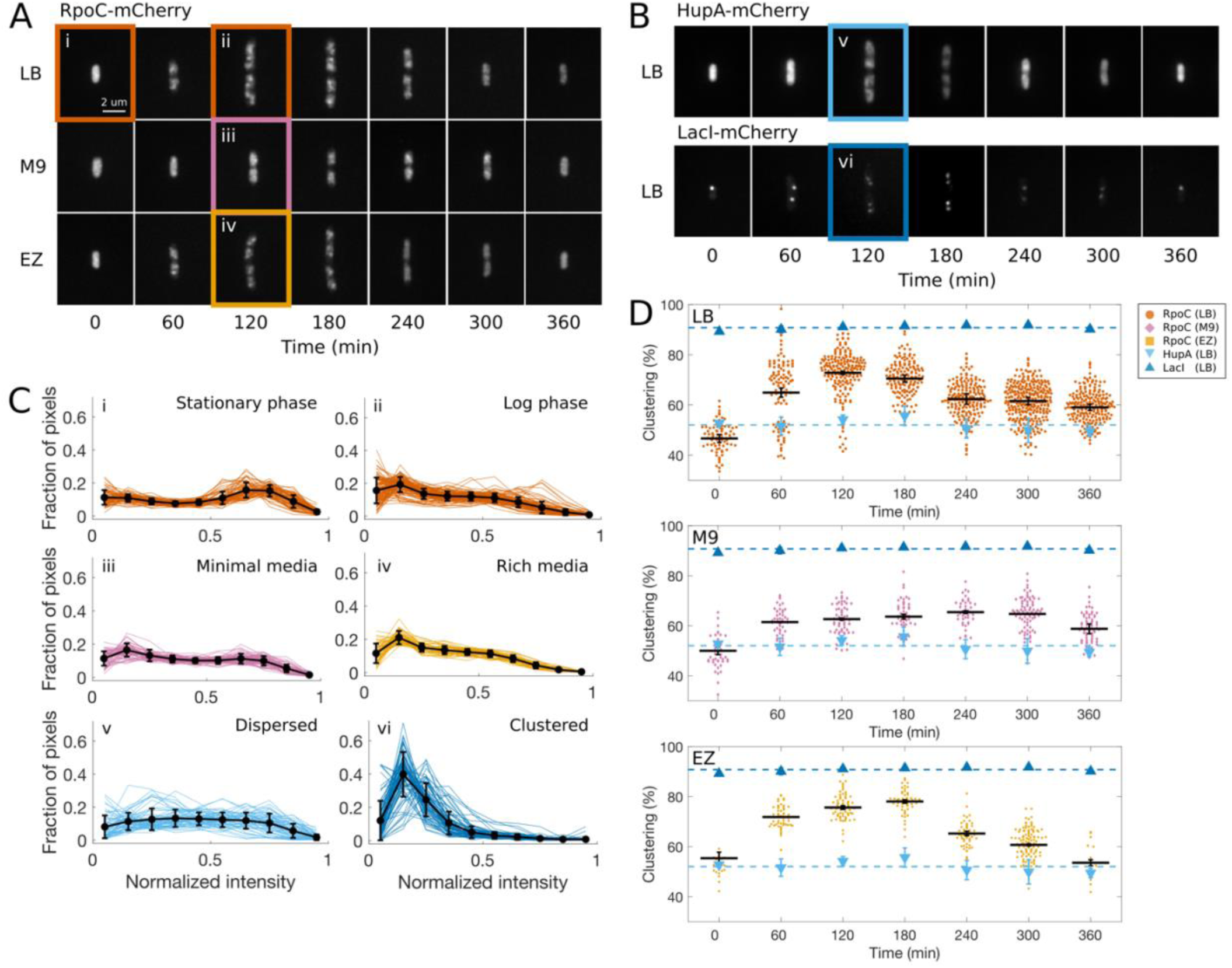
A) Fluorescence images of fixed cells expressing RpoC-mCherry collected during outgrowth at 37°C in rich (LB), minimal (M9) or defined (EZ) media. B) Fluorescence images of fixed cells expressing HupA-mCherry or LacI-mCherry collected during outgrowth at 37°C in LB. C) Normalized pixel intensity histograms for conditions highlighted in panels A and B (i-vi). D) Clustering of RpoC during outgrowth in LB (n = 6), M9 (n = 3) and EZ (n = 3). Data points correspond to individual cells; black bars represent means, error bars are standard error of the mean across n biological replicates. Triangles represent mean ± standard error of the mean for HupA and LacI controls (n = 3 each). Dashed lines are the means for HupA and LacI across all time points.

In fast growth conditions, the *E. coli* chromosome undergoes overlapping rounds of replication (38) and folds into a compact filamentous structure (39-41). Since RNAP binds to DNA, the observed clustering may result indirectly from changes in nucleoid morphology during outgrowth. To test this hypothesis, we repeated the outgrowth experiment with a strain expressing an mCherry fusion of HupA (42), a histone-like protein that binds throughout the nucleoid (43). Although the nucleoid becomes more dense in log phase (Fig. 1B, 120-180 min), we did not observe any distinct foci, suggesting that RNAP clustering is not caused by DNA compaction.

To quantify the spatial distribution of RNAP over time, we adapted a method previously established to monitor chromosome condensation (44) and nucleolar morphology (45). Briefly, individual cells were segmented from bright field images and the fluorescence intensity per pixel (I) was normalized by the minimum and maximum values in the segmented area, such that I_n_ = (I-I_min_)/(I_max_-I_min_). The resulting normalized pixel intensity histogram shifts from an approximately uniform distribution at 0 min (Fig. 1C, i) to a right-skewed distribution at 120 min (Fig. 1C, ii). We observed a similar shift from uniform to skewed distribution for log-phase cells in M9 compared to EZ (Fig. 1C, iii, iv). Histogram shape reflects the degree to which RNAP is dispersed or clustered in the cell. For comparison, HupA-mCherry provided a baseline for a dispersed localization pattern (Fig. 1C, v), while LacI-mCherry served as a control for a clustered pattern (Fig. 1C, vi). The latter strain contains a *lacO* array integrated near the terminus of the chromosome, resulting in one to four copies to which LacI-mCherry binds (Fig. 1B).

To compare histograms between individual cells and across different time points and media, we calculated the percent of pixels with a normalized intensity below a threshold, I_n_ < 0.5, for each cell. This percent corresponds to the area segmented as background, which increases upon protein clustering (Fig. S1). In LB, lag-phase cells have 47 ± 2% pixels below the threshold, while log-phase cells have 73 ± 1%, reflecting the transition from a dispersed to clustered organization (Fig. 1C, D). Interestingly, this transition appears to be discontinuous. At 60 min, the population is bimodal, with a majority of cells displaying clusters while 30 ± 7% of cells retain the dispersed localization pattern. By 120 min, nearly all cells have assembled clusters (Fig. 1D, LB). The discrete transition from dispersed to clustered organization is reminiscent of a first-order phase transition. RNAP also clusters in log-phase cells in M9, though to a lesser extent, and in EZ. The kinetics of clustering differ slightly between LB and EZ (Fig. 1D), perhaps due to the distinct nutrient composition of each medium.

### RNAP clustering is mediated by protein-protein interactions, not solely DNA-binding

Several mechanisms have been proposed to explain RNAP clustering. The most popular is the DNA-binding, or transcription factory, model (Fig. 2A). In this model, RNAP clusters are composed of polymerases that are directly bound to DNA and actively transcribing (31, 32). Under fast growth conditions, RNAP is densely packed on *rrn* operons (46) and the vast majority are engaged in synthesis of ribosomal RNA (rRNA) (47), prompting comparisons to the eukaryotic nucleolus (37). However, recent evidence indicates that nucleoid structure may play a more dominant role in RNAP clustering than transcriptional activity itself (36). Alternatively, we hypothesize that RNAP clusters may instead assemble primarily by LLPS such that RNAP is maintained at a high local concentration through weak multivalent protein-protein interactions, independent of its transcriptional activity (Fig. 2A).

**Figure 2.**
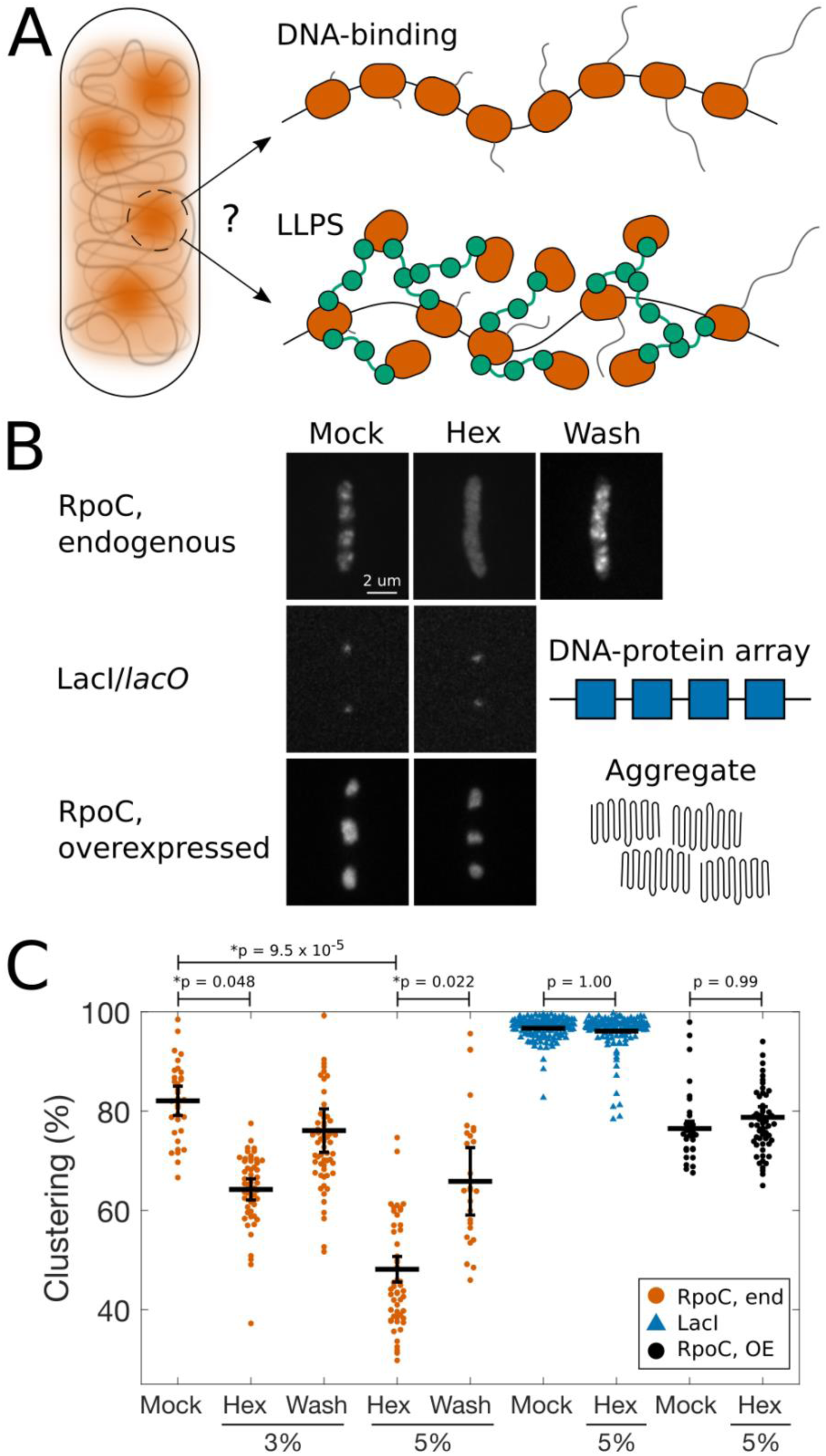
A) Possible mechanisms for RNAP clustering. The DNA-binding hypothesis proposes that RNAP molecules cluster through direct binding to DNA. Alternatively, RNAP molecules may instead cluster through liquid-liquid phase separation (LLPS). B) Fluorescence images of fixed cells expressing RpoC-mCherry from its native promoter (endogenous); LacI-mCherry; or RpoC-GFP from an IPTG-inducible promoter on a high copy number plasmid. Cells were treated with media (Mock) or 5% 1,6-hexanediol (Hex), and subsequently washed with fresh media to remove hexanediol (Wash). C) Quantification of clustering of fluorescent proteins for each treatment condition. Data points correspond to individual cells; black bars represent means, error bars are standard error of the mean across n = 3 biological replicates. *p* values calculated by ANOVA and Tukey-Kramer post hoc test.

To distinguish between these models, we treated cells with the aliphatic alcohol 1,6-hexanediol. This chemical dissolves liquid-like condensates, but not solid-like aggregates (48, 49) and has become a useful - though not definitive - assay for LLPS (50). Consistent with our hypothesis, RNAP clusters rapidly dispersed following addition of 1,6-hexanediol to cells grown in LB at 37°C for 120 min (Fig. 2B, C). The effect was dose-dependent and reversible, as RNAP clusters quickly reassembled when the hexanediol was washed out. Importantly, DNA-bound foci of LacI-mCherry are resistant to 1,6-hexanediol (Fig. 2B, C), demonstrating that this treatment does not interfere with direct binding to DNA. This observation is consistent with results from human cells, in which hexanediol does not disrupt nonspecific DNA binding of RNA polymerase II (51).

Finally, to determine whether RNAP clusters are physically distinct from protein aggregates, we used an IPTG-inducible plasmid to overexpress GFP-RpoC (52). When overexpressed, RpoC accumulates in the cytoplasm, rather than the nucleoid, and forms large aggregates at mid-cell and the poles. As expected, these aggregates are not affected by hexanediol treatment (Fig. 2B, C). Together, these results suggest that weak protein-protein interactions contribute to RNAP clustering and that it is not mediated solely by DNA-binding.

### Antitermination factors are required for RNAP clustering

Next, we sought to identify molecular components that are required for RNAP clustering. We began by compiling a list of proteins that contain disordered regions. Among the most highly-expressed disordered proteins in *E. coli* are RpoZ, an RNAP subunit; NusA and NusG, antitermination factors that interact directly with RNAP and each other; and H-NS, a nucleoid-associated protein which negatively regulates *rrn* transcription (53). To identify additional candidates, we calculated the mean net charge and mean hydrophobicity for all 4351 proteins in the *E. coli* proteome (Fig. 3A). Previous sequence analyses identified an empirical relationship that distinguishes between folded and natively unfolded proteins based on their amino acid composition (54). Using this classifier, we found 202 proteins, or 4.6% of the proteome, that are likely unfolded. We focused on proteins that are known to be involved in rRNA transcription (Table 1). Of this set, five proteins, or 25%, fall above the unfolded/folded boundary on the charge-hydrophobicity plot (Fig. 3A, Table 1). Notably, RpoZ, NusA and NusG were not identified by this method. Finally, disordered regions in several of these proteins have been confirmed experimentally (55-61). Taken together, transcriptional regulators in *E. coli* appear to be enriched for disordered proteins, as they are in eukaryotes (25, 62).

**Table 1.** Proteins involved in rRNA transcription. Proteins with evidence for disorder are highlighted in **bold**.

| RNA polymerase | Transcription factors | Antitermination factors | Ribosomal proteins | Regulators |
| --- | --- | --- | --- | --- |
| RpoA | **DksA**<sub>x+</sub>* | **NusA**<sub>x+</sub> | **S4**<sub>x+</sub>* | RelA |
| RpoB | **Fis**<sub>+</sub> | NusB | S10/NusE | SpoT |
| RpoC | **H-NS**<sub>x+</sub>* | NusE/S10 | L3 |  |
| **RpoD**<sub>x</sub>* | Lrp | **NusG**<sub>x+</sub> | **L4**<sub>+</sub> |  |
| **RpoZ**<sub>x</sub> |  | SuhB | **L13**<sub>+</sub>* |  |

Evidence for disorder: <sub>x</sub>Predicted; <sub>+</sub>Experimentally confirmed; \*Figure 3A

**Figure 3.**
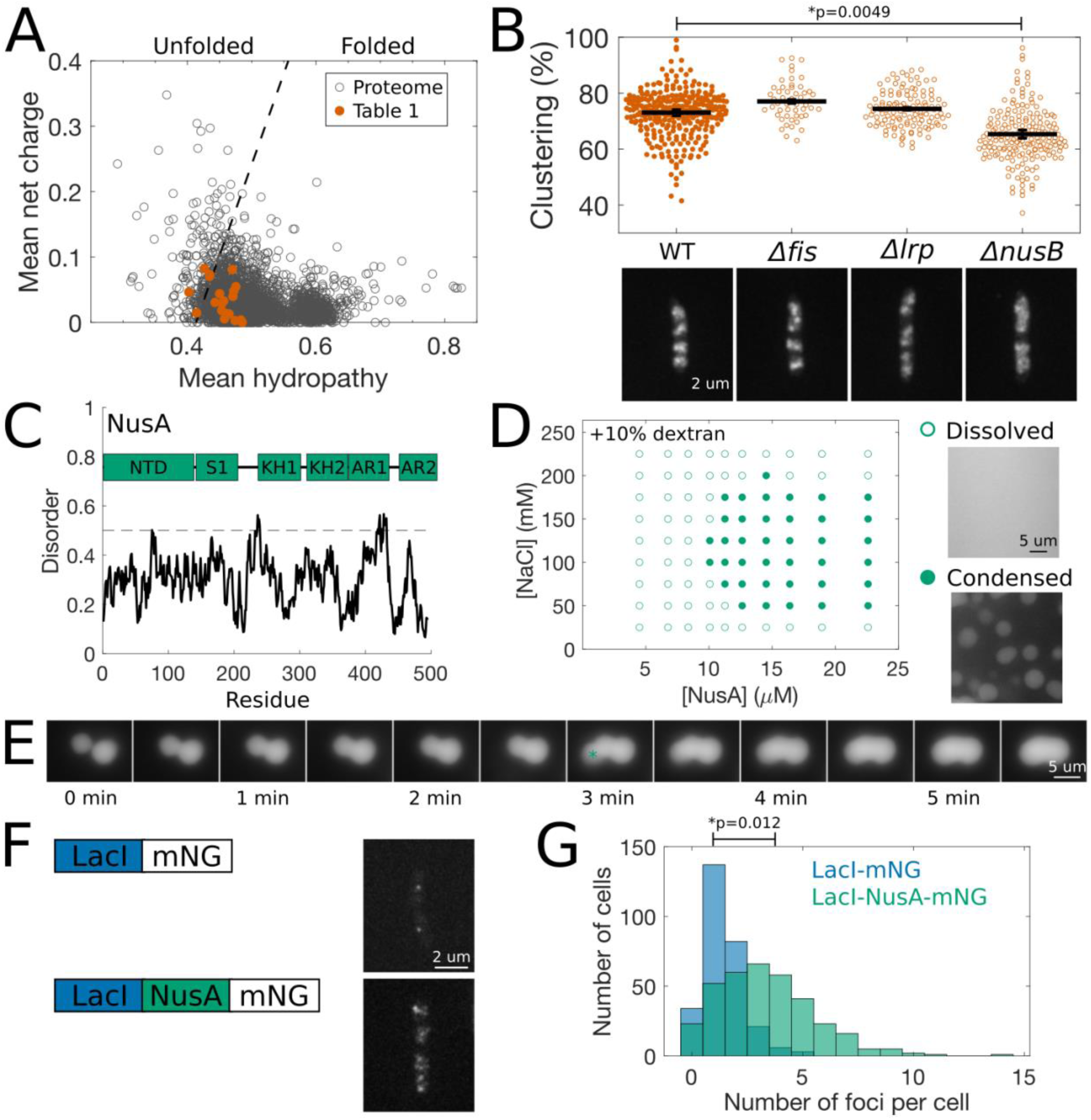
A) Charge v. hydropathy plot for the *E. coli* proteome, with proteins from Table 1 highlighted. The dashed line represents an empirical border between proteins that are natively unfolded (left) and folded (right). B) Clustering and fluorescence images of wildtype and deletion mutants expressing RpoC-mCherry. *p* values were calculated by ANOVA and Tukey-Kramer post hoc test. C) Domain structure and predicted disorder (by IUPred) of NusA. D) Phase diagram for purified NusA in the presence of 10% dextran. Open circles indicate conditions in which the protein is dissolved; closed circles indicate conditions in which the protein is condensed. E) Montage of NusA droplet fusion. A third droplet falls from solution at t = 3 min (marked by *). F) Fluorescence images of live cells expressing LacI-mNeonGreen or LacI-NusA-mNeonGreen. G) Histogram of number of foci per cell for each protein construct (n = 3). *p* value was calculated by unpaired, two-tailed Student’s t test.

To determine whether candidate proteins contribute to RNAP clustering, we transduced the rpoC-mCherry marker into available mutants from the Keio knockout collection (63). The resulting strains were grown in LB at 37°C and cells were collected at 120 min, fixed with formaldehyde and imaged. First, we examined the spatial organization of RNAP in the absence of Fis, a transcriptional activator (64), and Lrp, a transcriptional repressor (65). Fis is a strong candidate for LLPS: its N-terminal domain is disordered (57) (Fig. S2B); it has multiple binding sites upstream of each *rrn* promoter (66); and it is highly expressed during log phase (67), when RNAP clusters assemble (Fig. 1). Moreover, we found that purified GFP-Fis condenses into liquid-like droplets in a salt-dependent manner *in vitro* (Fig. S3A). Unexpectedly, however, deletion of Fis does not impair RNAP clustering, as RpoC-mCherry still assembled into bright foci in *Δfis* cells (Fig. 3B).

Lrp contains a flexible linker that connects folded N- and C-terminal domains (68), but is otherwise ordered (Table 1, Fig. S2C). Therefore, this protein is not expected to phase separate and serves as a negative control. We observed no change in RNAP clustering in *Δlrp* cells compared to wildtype (WT) (Fig. 3B).

NusA and NusG are essential in *E. coli* (69), so we next tested NusB for its role in RNAP clustering. NusB colocalizes with RNAP clusters (70) and, together with NusA, NusG, NusE (the 30S ribosomal protein S10) and SuhB, forms a complex that prevents premature Rho-dependent termination of *rrn* genes (71, 72). When we visualized RpoC-mCherry in the *ΔnusB* strain, we observed a more dispersed localization pattern (Fig. 3B). RNAP is distributed throughout the nucleoid of *ΔnusB* cells and concentrates into just a few dim foci. This result suggests that NusB, or another antitermination factor (Table 1), is required for efficient RNAP clustering.

### NusA undergoes LLPS *in vitro*

NusA is an abundant protein composed of six folded domains connected by flexible linkers (Fig. 3C). The central S1 motif and two K homology (KH) domains bind RNA (73, 74), while the N-terminal domain (NTD) and two C-terminal acidic repeat (AR) domains interact with a variety of proteins, including multiple subunits of RNAP (75-77), NusG (78), SuhB (71, 72) and the bacteriophage lambda protein N (79, 80). AR2 can also interact with KH1 to block RNA binding (76, 81). This modular architecture, which facilitates both heterotypic and homotypic interactions, is consistent with the recent “stickers-and-spacers” model used to describe LLPS (82).

To test its intrinsic ability to phase separate, we purified GFP-NusA and monitored its behavior *in vitro* using complementary biochemical assays (83, 84). First, we looked for condensed-phase protein droplets by fluorescence microscopy. Second, we measured the protein concentration of NusA solutions following centrifugation. In this latter assay, the final concentration matches the initial concentration when NusA remains dissolved. However, if NusA phase separates, then centrifugation collects the dense phase in the pellet and the (lower) concentration of the supernatant corresponds to the saturation concentration at the phase boundary. We varied the salt concentration in our assay buffer to modulate the affinity of intermolecular interactions and map the phase diagram for NusA.

Initially, we failed to detect phase separation by either technique. NusA remains soluble up to at least 22.5 μM at all salt concentrations tested (25 - 225 mM NaCl) (Fig. S3B). To mimic the crowded environment inside a cell, we repeated these assays in the presence of 10% dextran, which has been used to induce phase separation of both prokaryotic and eukaryotic proteins (85, 86). Under these conditions, we consistently observed phase separation and found qualitative and quantitative agreement between the two assays (Fig. 3D). In the presence of dextran, GFP-NusA forms large spherical droplets at physiological salt concentrations above a saturation concentration of ∼10 μM. Time-lapse imaging revealed that multiple droplets fuse upon contact, demonstrating liquid-like properties (Fig. 3E). Consistent with the dispersal of clusters by hexanediol *in vivo* (Fig. 2), we also found that NusA droplets dissolved in 10% hexanediol *in vitro* (Fig. S3C). These results indicate that NusA is capable of forming homotypic interactions that can drive LLPS. However, as we found for Fis (Fig. S3A), they do not necessarily imply that NusA phase separates *in vivo*.

### NusA nucleates foci *in vivo*

To determine whether NusA can nucleate condensates in cells, we used the *lacO*/LacI system in a strategy similar to that recently applied to eukaryotic transcription factors (87). We generated two constructs: (1) LacI-mNeonGreen alone and (2) LacI-NusA-mNeonGreen, which contains the full-length NusA protein; expressed them in cells containing a *lacO* array near the terminus of the chromosome; and counted the number of fluorescent foci in each cell (Fig. 3F). Cells expressing LacI-mNG had 1.4 ± 0.2 foci per cell (Fig. 3G), consistent with the copy number of the *lacO* array (88). In contrast, cells expressing the NusA fusion protein (LacI-NusA-mNG) had significantly more foci, with 3.9 ± 0.6 per cell (Fig. 3F, G). Since the number of fluorescent foci exceeds the number of *lacO* arrays, this result suggests that NusA phase separates *in vivo*, independently of specific DNA binding.

### Components of RNAP clusters are dynamic

A defining feature of liquid-like condensates is the dynamic behavior of their components, which rearrange within the condensate and exchange with the surrounding bulk phase (15). To investigate the dynamics of RNAP clusters, we used single-molecule tracking. We generated new strains expressing either RpoC or NusA fused to the photoconvertible protein mMaple3 (89). Upon activation with 405-nm light, mMaple3 switches from the green channel to the red, allowing us to visualize single molecules over time (Fig. 4A). Using photoactivated localization microscopy (90), we acquired time-lapse movies of live cells growing in EZ medium at 37°C and tracked the position of RpoC and NusA molecules at 20-ms intervals.

**Figure 4.**
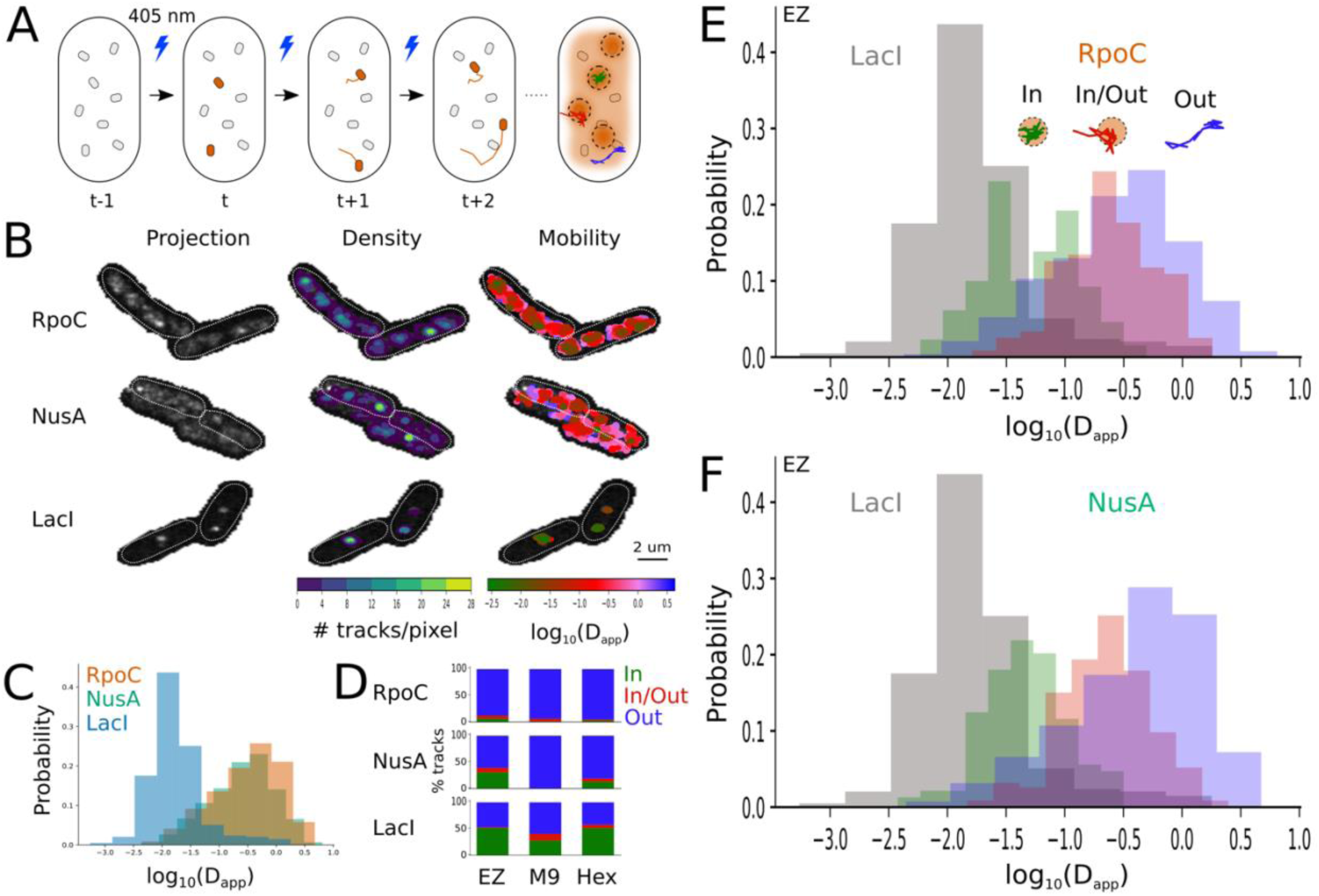
A) Schematic of a single-molecule tracking experiment. Cells expressing mMaple3 fusion proteins are continuously activated with 405-nm light and converted molecules are tracked over time. B) Maximum intensity projections of pairs of cells expressing RpoC-mMaple3, NusA-mMaple3 or LacI-mMaple3. The density and mobility, respectively, of tracks are overlaid. C) Distribution of D_app_ for all tracks. D) Classification of tracks for each protein under three growth conditions. “In” tracks are localized to clusters; “In/Out” tracks exchange between clusters and the bulk nucleoid; “Out” tracks do not interact with clusters. E) Distribution of D_app_ for each class of RpoC tracks. F) Distribution of D_app_ for each class of NusA tracks.

For both proteins, we observed tracks throughout the nucleoid, with a high density of tracks localized to two to four clusters per cell (Fig. 4B). To quantify the mobility of these molecules, we calculated the apparent diffusion coefficient (D_app_) of each track. Both RpoC and NusA gave broad distributions, ranging from 5 × 10^−3^ to 5 μm^2^/s, with an average D_app_ of 0.52 ± 0.01 μm^2^/s and 0.63 ± 0.01 μm^2^/s, respectively (Fig. 4C, Table 2). This wide range of mobilities is similar to previous estimates for RpoC (34, 91) and likely reflects different states of activity, for example molecules engaged in transcription or molecules non-specifically bound to DNA (91). Interestingly, there appeared to be a correlation between track density and mobility, with molecules in high-density clusters tending to move more slowly than molecules in low-density regions of the nucleoid (Fig. 4B).

**Table 2.** Apparent diffusion coefficients (μm^2^/s) from single-molecule tracking. Values are mean ± standard error across n = 4-6 biological replicates.

| Protein | Class | EZ | M9 | Hex |
| --- | --- | --- | --- | --- |
| RpoC | ALL | $0.52 \pm 0.10$ | $0.72 \pm 0.04$ | $0.55 \pm 0.03$ |
| | In | $0.16 \pm 0.04$ | $0.37 \pm 0.05$ | $0.26 \pm 0.01$ |
| | In/Out | $0.44 \pm 0.04$ | $0.643 \pm 0.003$ | $0.57 \pm 0.02$ |
| | Out | $0.61 \pm 0.13$ | $0.81 \pm 0.02$ | $0.66 \pm 0.04$ |
| NusA | ALL | $0.63 \pm 0.01$ | $0.77 \pm 0.02$ | $0.57 \pm 0.04$ |
| | In | $0.18 \pm 0.09$ | $0.097 \pm 0.003$ | $0.056 \pm 0.002$ |
| | In/Out | $0.47 \pm 0.12$ | $0.44 \pm 0.03$ | $0.40 \pm 0.02$ |
| | Out | $0.76 \pm 0.02$ | $0.85 \pm 0.06$ | $0.73 \pm 0.05$ |
| LacI | ALL | $0.11 \pm 0.03$ | $0.11 \pm 0.02$ | $0.11 \pm 0.01$ |
| | In | $0.023 \pm 0.004$ | $0.023 \pm 0.002$ | $0.021 \pm 0.002$ |
| | In/Out | $0.65 \pm 0.07$ | $0.56 \pm 0.05$ | $0.22 \pm 0.04$ |
| | Out | $0.37 \pm 0.17$ | $0.32 \pm 0.04$ | $0.37 \pm 0.06$ |

If clustering were caused by DNA binding, then RNAP molecules in a cluster would have the same mobility as a DNA locus. However, if clustering were instead mediated by LLPS, then RNAP molecules may move differently inside a cluster compared to when actively transcribing DNA or diffusing through the rest of the nucleoid. To measure the baseline mobility of a DNA locus, we tracked LacI-mMaple3 molecules in a strain carrying a *lacO* array (90). We observed tracks in only one or two locations per cell, suggesting that most LacI molecules are stably bound to the DNA array (Fig. 4B). Consistent with this interpretation, and with previous measurements of DNA motion in *E. coli* (92), the D_app_ distribution was much narrower for LacI, with a peak near 1 × 10^−2^ μm^2^/s (Fig. 4C). Moreover, the LacI distribution is shifted left relative to the RpoC and NusA distributions, indicating that the majority of RpoC and NusA molecules move faster than DNA. Nevertheless, there is substantial overlap among the distributions (Fig. 4C). The slow-moving RpoC molecules in this overlap region are likely engaged in transcription, while the NusA molecules are engaged in co-transcriptional antitermination. This population of molecules could correspond to all clustered molecules, in support of the DNA-binding model, or only a subset of clustered molecules, in support of the LLPS model (Fig. 2A).

To distinguish between these possibilities, and to compare the mobility of molecules in RNAP clusters to the mobility of DNA directly, we developed a stringent classification scheme. First, we created maximum intensity projections over time to identify regions with a high density of activated molecules (Fig. 4B). Second, we defined a “cluster” as any region detected in the projection that overlapped with a minimum of three tracks. Finally, a molecule was designated as “In”, if its entire trajectory was contained inside a cluster; “In/Out”, if its trajectory crossed a cluster boundary but spent a majority of timepoints inside a cluster; and “Out”, if its trajectory never overlapped with a cluster. This classification scheme is conservative. We intentionally set a strict definition for the “In” class, preferring to underestimate the number of “In” molecules in order to calculate their mobility more accurately. As a result, only ∼5-10% of RpoC tracks and 10-30% of NusA tracks were assigned to the “In” class (Fig. 4D). Consequently, the “Out” class likely contains many false positives, as reflected by the large percentage of molecules assigned to this class (Fig. 4D), even for LacI, and by the leftward skew of the D_app_ distribution for this class (Fig. 4E, F).

To validate our classification scheme, we applied two independent methods to the raw (unclassified) data (Fig. 4C). First, we used a nonparametric Bayesian approach called single-molecule analysis by unsupervised Gibbs sampling (SMAUG) (93). This method identifies the number of different mobility states in a population of trajectories and calculates the fraction of molecules and apparent diffusion coefficient in each state. Second, we fit a gaussian mixture model to the full distribution of D_app_ for all tracks. In both cases, we identified three to four sub-populations of molecules with mobilities that are consistent with our results from single-molecule tracking (Fig. S4).

After classifying tracks from each strain (Fig. 4D), we compared the normalized distributions of D_app_ for each class. RpoC molecules classified as “In” clusters had a bimodal distribution, with the lower peak overlapping the distribution for LacI (Fig. 4E). The higher peak was shifted to the right, centered at ∼0.1 μm^2^/s, which is still an order of magnitude smaller than the majority of RpoC molecules. These results suggest that RNAP clusters contain two populations of RpoC: one that is bound to DNA and another that is not bound, moves more quickly and yet remains localized in high-density clusters. We propose that the former population is engaged in transcription, while the latter population is diffusing within a surrounding liquid-like condensate. In addition, the distribution for “In/Out” molecules overlapped with this high-mobility “In” population and also extended to much greater values of D_app_. This is consistent with molecules that are exchanging between a condensed phase and the bulk nucleoid, and which spend variable amounts of time in each phase. Finally, the “Out” distribution was shifted farthest right, to a mean of 0.61 ± 0.1 μm^2^/s (Table 2). The high mobility of these molecules suggests that they are diffusing freely through the cytoplasm and/or non-specifically bound to the nucleoid. The long left tail of this distribution likely arises from the misclassification of tracks whose clusters were not detected. Alternatively, some molecules in this tail may be engaged in transcription at other sites throughout the nucleoid.

NusA exhibited similar dynamics (Table 2). Like RpoC, the D_app_ distribution for the “In” class of NusA partially overlapped with LacI (Fig. 4F), indicating that some NusA molecules are likely associated with DNA (presumably indirectly through interactions with RNAP (75, 76)). Importantly, this class also contains faster-moving molecules, consistent with the LLPS hypothesis. The “In/Out” and “Out” distributions are shifted progressively to the right, with mean values comparable to RpoC (Table 2). These data reveal that, like RpoC, NusA exists in several different states in the cell, such that its mobility varies both in space and with molecular density.

Finally, we determined how the dynamics of RpoC and NusA change under different growth conditions. We repeated single-molecule tracking in live cells grown in M9 or in EZ after treatment with 5% 1,6-hexanediol (Hex). As expected, the percent of tracks that were classified as “In” clusters decreased in both conditions (Fig. 4D), consistent with the reduced RNAP clustering seen in fixed cells (Fig. 1A, E and Fig. 2B, C), as well as the dissolution of NusA droplets in the presence of hexanediol *in vitro* (Fig. S3). Furthermore, while the mobility of DNA-bound LacI did not change, the few RpoC molecules that remained in clusters appeared less constrained (Table 2, Fig. S5). This may be due to changes in the composition or molecular density of the condensates, which could affect their viscosity. Conversely, the mobility of NusA molecules in clusters became more restricted, approaching the motion of a DNA locus. Together, these results suggest that when clusters dissolve in M9 or Hex, NusA disperses into the cytoplasm while RNAP stays localized through transient and non-specific binding to DNA.

## Discussion

Our results demonstrate that RNAP clusters are biomolecular condensates that assemble through LLPS. We found that clustering occurs rapidly during outgrowth in rich media, when cells transition from lag phase to log phase (Fig. 1). We showed that RNAP clusters, but not protein bound to DNA arrays or protein aggregates, dissolve upon treatment with hexanediol (Fig. 2). This sensitivity is dose-dependent and reversible, suggesting that hexanediol acts as a solvent to disrupt the intermolecular interactions driving LLPS. We identified NusA as a potential mediator of these interactions (Fig. 3), though additional components may be involved. Deletion of NusB, a component of the antitermination complex, perturbs clustering in cells, while NusA phase separates *in vitro* and nucleates foci *in vivo*. Finally, we demonstrated that molecules inside RNAP clusters are dynamic, moving more quickly than DNA loci but more slowly than molecules in the bulk nucleoid (Fig. 4). Together, these results provide direct evidence for the LLPS hypothesis, establishing LLPS as a mechanism for intracellular organization in bacteria.

Our work has important implications for transcriptional regulation in particular and bacterial cell biology in general. It also highlights single-molecule tracking as a powerful tool for examining LLPS in bacteria and beyond.

### Partitioning of RNAP

Early biochemical work established that the macromolecular composition of *E. coli* cells varies with growth rate (94, 95). In particular, the total number of RNAPs increases with growth rate, and active RNAPs redistribute from protein-coding genes to stable RNA (i.e. rRNA and tRNA) genes as doubling time decreases (47). These data were used as support for the DNA-binding model when RNAP clusters were first observed by microscopy (31, 32). They also inspired the development of theoretical models to predict how RNAP is partitioned among various states, such as active transcription of mRNA or rRNA, non-specific binding to the nucleoid, or free diffusion in the cytoplasm (96, 97).

More recent single-molecule tracking studies assigned RNAP to different states based on its mobility (34, 91). For example, slow-moving molecules (D < 0.03 μm^2^/s) were classified as transcribing; fast-moving molecules (D = 0.7 μm^2^/s) as freely diffusing; and molecules with intermediate mobility (D = 0.21 μm^2^/s) as binding non-specifically to the nucleoid (91). The D_app_ values presented in Fig. 4 are comparable to these previous measurements, but our interpretation differs. Instead of assigning all slow-moving molecules to the active transcription state, we favor dividing this group of molecules into two distinct states and introducing an additional “condensed but non-transcribing” state.

Our partitioning scheme is not only consistent with the LLPS hypothesis, it also resolves a discrepancy in the literature. Numerical results (97) agree with gene expression data (98) but not with single-molecule tracking (91). Specifically, Klumpp and Hwa (97) predicted that ∼21% of RNAP is actively transcribing at a doubling time of 40 min, while Bakshi *et al.* (91) estimated 49%. If the updated number of transcribing RNAP were plugged back into Klumpp’s model, then it would overestimate Nomura’s expression data (98). However, if ∼30% of the slow-moving molecules that Bakshi classified as “transcribing” were not actually transcribing, then the numerical, expression and single-molecule results would be consistent. An alternative interpretation for this discrepancy is that the mRNA elongation rate is slower than assumed in the model.

### Transcriptional regulation

RNAP clusters assemble under fast growth conditions, where rRNA synthesis accounts for ∼80-90% of cellular transcription (47). This correlation between clustering and rRNA transcription prompted early comparisons to the eukaryotic nucleolus (31, 32). The nucleolus is a biomolecular condensate composed of immiscible liquid phases that spatiotemporally coordinate ribosome biogenesis at rDNA sites (16, 99-101). By analogy, bacterial RNAP clusters were thought to colocalize with *rrn* operons and function in rRNA synthesis. Indeed, RNAP clusters were recently shown to colocalize with nascent pre-rRNA, confirming that at least some RNAPs within clusters are actively transcribing rRNA (36). Surprisingly, however, this study also found that RNAP clusters persisted even when rRNA transcription was perturbed. These results contradict expectations for the DNA-binding model, but are fully consistent with an LLPS model in which phase separation is mediated primarily by protein-protein interactions, rather than RNA-protein interactions. Furthermore, the size of RNAP clusters is smaller after disruption of rRNA transcription (36). This behavior is similar to the nucleolus, which is stabilized by rRNA, but whose protein components nevertheless condense into small droplets in the absence of rRNA transcription in early *C. elegans* embryos (17).

Ribosome content is limiting for growth, such that bacteria must allocate considerable cellular resources toward ribosome synthesis (102). Thus, it may at first seem wasteful for cells to partition a large fraction of RNAP into a condensed, non-transcribing state. However, a simple calculation (see below) suggests that this state does not compete with the actively transcribing state. In fact, the condensed pool of RNAPs may serve to accelerate re-initiation rates in order to maintain a maximal level of rRNA transcription. *E. coli* has seven *rrn* operons, with four positioned near the origin of replication, such that there are up to 50 copies per cell under optimal growth conditions (37). Each operon is 5.5 kb long, with RNAPs spaced every ∼85 bp (46). Altogether, if every *rrn* is fully saturated with RNAPs, then 50 *rrn*/cell x 5.5 kb/*rrn* x 1 RNAP/85 bp = 3235 RNAPs are expected to be in the transcribing state. Yet this fraction represents just one-third of the ∼10,000 RNAPs (47), leaving more than enough to populate the condensed, mRNA-transcribing, non-specifically bound and freely-diffusing states. Furthermore, these estimates are consistent with Klumpp’s numerical model (97), as well as previous single-molecule studies. PALM analyses conducted at slower growth rates (31 - 73 min doubling times) counted between 70 and 500 molecules per cluster and 2 to 4 clusters per cell (33, 34, 36), out of a total of ∼5000 RNAPs per cell. Therefore, at both fast and intermediate growth conditions, less than half of all RNAPs are engaged in rRNA transcription and we propose that some of the non-transcribing pool is partitioned into condensates.

LLPS provides a versatile mechanism for cells to control the localization, accessibility and/or activity of macromolecules in response to internal and external cues (103, 104). Here, we have shown that *E. coli* harnesses LLPS to reorganize its transcriptional machinery in response to nutrient availability. Further investigations will be necessary to dissect how known transcriptional regulation pathways, including DksA and (p)ppGpp (105), affect the propensity of NusA and other molecular candidates to phase separate, and how clustering influences transcription and subsequent processing of rRNA (106).

### Biomolecular condensates in bacteria

In addition to RNAP clusters, bacteria likely contain a variety of biomolecular condensates. Many candidates have already been identified. For example, RNase E assembles into bacterial RNP bodies which are required for efficient mRNA turnover and confer tolerance to stress in *Caulobacter* (28). Several DEAD-box RNA helicases have been proposed to form condensates in *E. coli* (27, 107), but further investigations are necessary to determine whether they undergo LLPS at endogenous expression levels. Polyphosphate granules are another promising candidate. These spherical structures are synthesized during nitrogen starvation in *Pseudomonas* and appear to nucleate, grow and ripen like liquid droplets (108). In stationary phase, Dps compacts the *E. coli* nucleoid into a dense crystalline-like structure that excludes restriction enzymes yet permits access to RNAP (109). LLPS may also occur on the bacterial membrane, as the ABC transporter Rv1747 phase separates *in vitro* and forms clusters in *Mycobacterium* (110). There is also *in vitro* evidence that FtsZ and SlmA may form condensates that regulate the positioning of Z-ring assembly during *E. coli* cell division (85). Together, this growing list of putative condensates suggests that LLPS is ubiquitous in bacteria. These examples span many distantly related species; they appear in all phases of growth (i.e. lag, log, stationary); and they are associated with diverse cellular processes, from gene expression and metabolism to dormancy.

### Single-molecule tracking as a tool for examining LLPS *in vivo*

Despite this exciting progress, it remains challenging to study phase separation in any organism but particularly in bacteria. Indeed, while the appearance of fluorescent foci in cells is promising, this alone is ***not*** compelling evidence for LLPS because foci can form through many alternative mechanisms (51). So additional tools for interrogating putative condensates are urgently needed. Fluorescence recovery after photobleaching (FRAP) is one of the most commonly used tools, as it can probe both the dynamic rearrangement of molecules within a condensed phase and their rapid exchange with the surrounding environment (14). However, concerns about the liberal interpretation of this method have recently been raised (29). Furthermore, FRAP cannot easily be applied to the submicron structures in bacteria, nor can shape or fusion analysis (99).

Fortunately, single-molecule tracking has recently emerged as a powerful tool for examining LLPS *in vivo* (51, 111, 112). Indeed, the strongest evidence for bacterial condensates to date comes from single-molecule tracking of proteins in polar microdomains in *Caulobacter* (113). This study revealed that PopZ imposes a diffusion barrier to cytosolic proteins and restricts the mobility of signaling molecules at the cell poles. Our analysis of RNAP mobility (Fig. 4) further underscores the promise of applying single-molecule tracking to characterize the physical properties and functional consequences of biomolecular condensates in bacteria.

In summary, we have identified a novel biomolecular condensate in *E. coli*. Future work will clarify the molecular components and interactions that drive assembly of RNAP clusters, and determine how they affect gene expression, ribosome biogenesis and cell growth and size.

## Materials and Methods

### Bacterial strains

Strains used in this study are listed in Table 3. New strains were constructed by P1 transcription (114) or lambda red recombination (115), as appropriate.

**Table 3.**
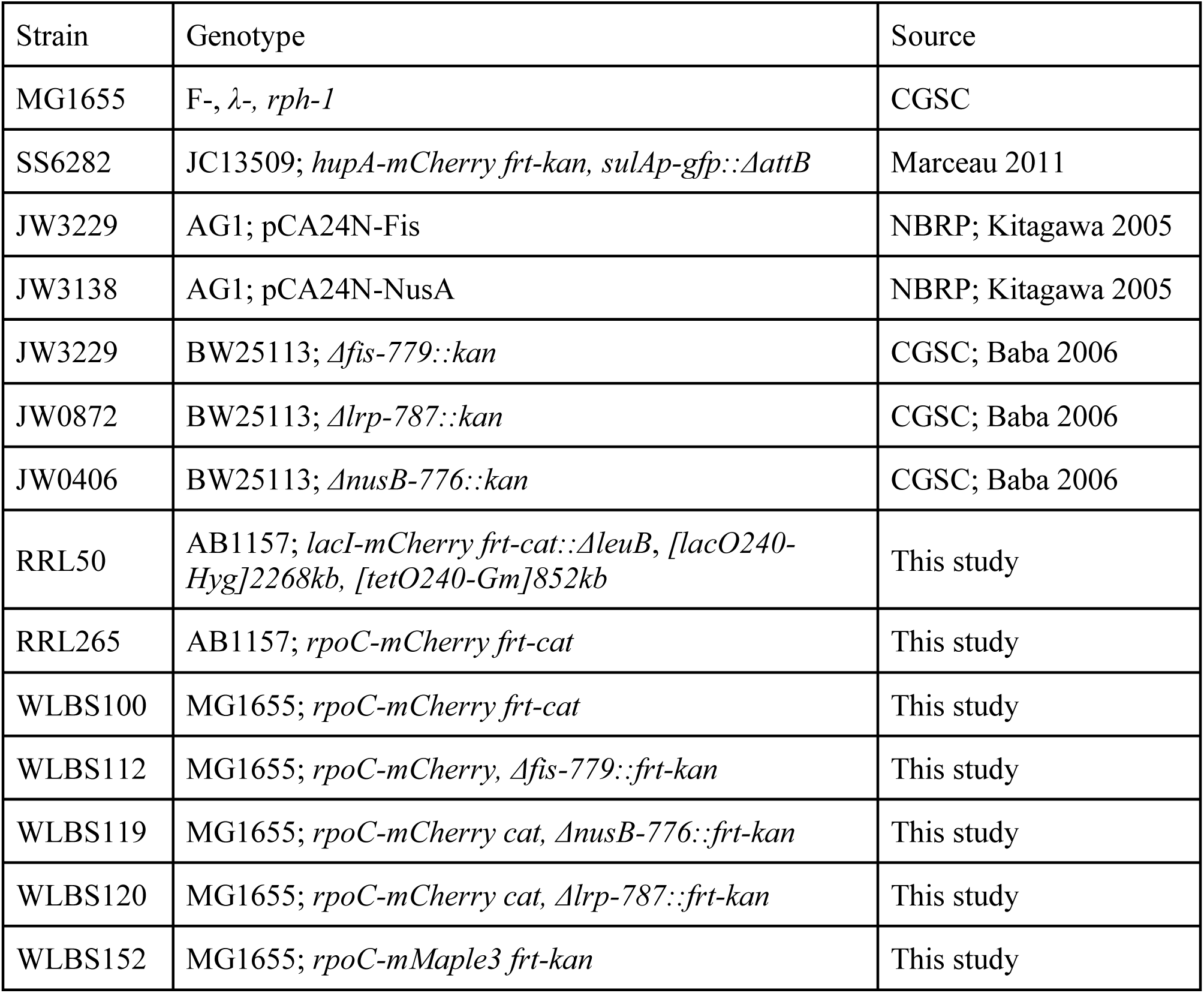
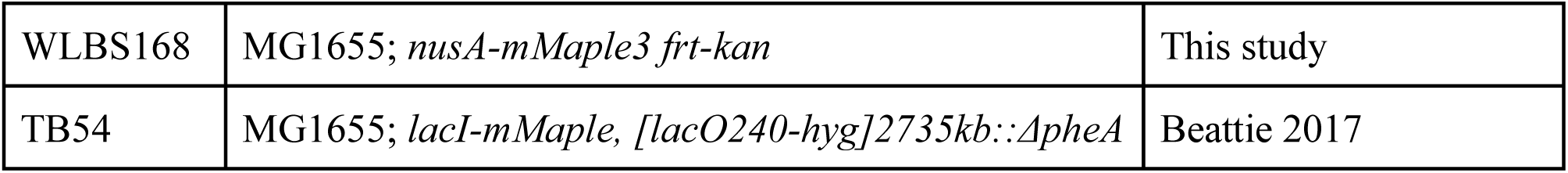
Genotypes of strains used in this study.

**Table 4.**
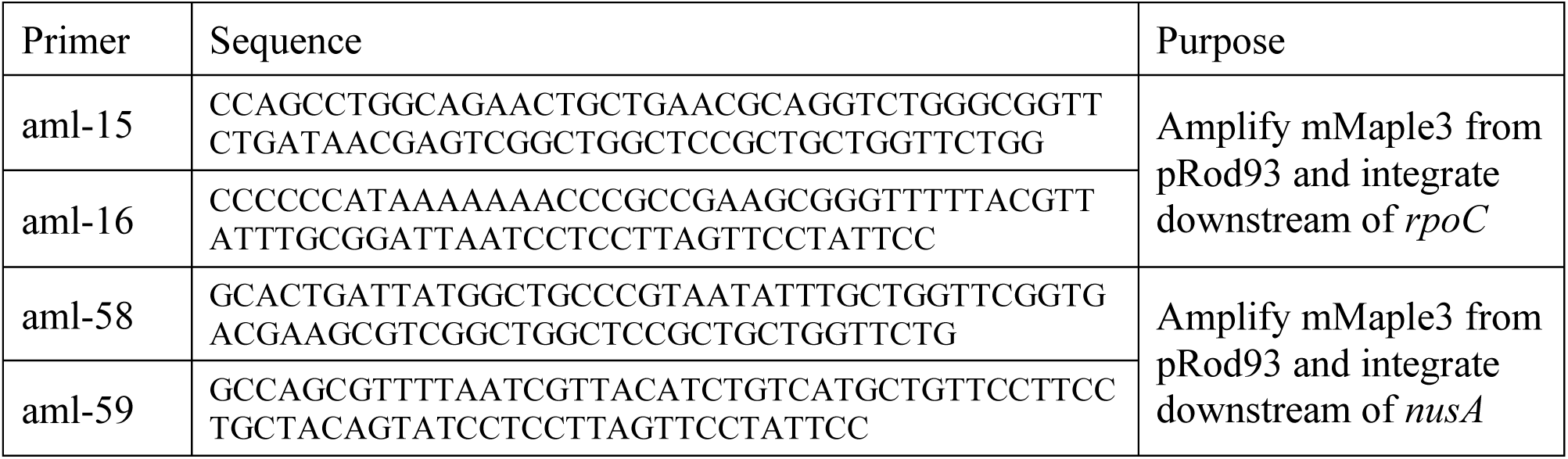
Oligonucleotides used in this study.

**Table 5.** Plasmids used in this study.

| Plasmid | Features | Source |
| --- | --- | --- |
| pKD46 | Lambda red genes | Datsenko 2000 |
| pROD93 | mMaple Kan R6K gamma ori, for C-terminal insertions | Beattie 2017 |

WLBS100: P1 transduction from RRL265 into MG1655

WLBS112: P1 transduction from JW3229 into WLBS100

WLBS119: P1 transduction from JW0406 into WLBS100

WLBS120: P1 transduction from JW0872 into WLBS100

WLBS152: PCR from pROD93 using primers aml-15, 16; Lambda red into MG1655/pKD46

WLBS168: PCR from pROD93 using primers aml-58, 59; Lambda red into MG1655/pKD46

### Growth conditions

Liquid cultures were inoculated from a single colony and grown over night at 37°C in LB. Cultures were diluted into fresh LB to an OD600 of ∼0.01 and grown for another 16 hrs. Cells were diluted again into fresh medium (LB, M9 or EZ), as indicated, for imaging. M9 minimal medium and EZ Rich Defined Medium (Teknova) were supplemented with 0.2% (w/v) glucose.

### Time-course imaging of fixed cells

Samples were collected every 60 min; fixed in 4% formaldehyde with constant mixing for 30 min at 37°C; washed three times in M9 at room temperature; and stored at 4°C. Fixed cells were mounted on 1% M9-agarose pads and imaged on an inverted Leica DMI 6000B equipped with a 100X 1.46 NA objective lens, a spinning disk confocal head (Yokogawa CSU10) and an EM-CCD camera (Hamamatsu ImagEM). Five to ten fields of view were randomly acquired for each time point.

Image analysis was done using custom scripts in Matlab (Mathworks). Cells were segmented from bright field images and fluorescence intensity values of all pixels within a cell were extracted. Clustering was calculated as described in Maddox 2006.

### Protein purification

6xHis-tagged GFP-Fis and GFP-NusA were expressed from pCA24N and purified using Ni-NTA resin (Qiagen) in a gravity-flow column at 4°C as described (Kitagawa 2005). Proteins were stored in 300 mM NaCl at −80°C and buffer-exchanged in spin columns (Amicon) just prior to condensation assays.

### Single-molecule imaging in live cells

Cells were grown in fresh medium (EZ or M9) for 90 min before mounting on 1% agarose pads, made with the same media. Pads were assembled in Gene Frames (1.0 x 1.0 cm, Thermo Fisher) with clean cover slips. Briefly, cover slips were soaked in Versa-Clean (Fisher) overnight; washed in methanol and acetone; sonicated for 30 min; baked in a plasma oven for 15 min; and flamed prior to mounting. Samples were maintained at 37°C throughout imaging.

Images were acquired on an inverted Olympus IX83 with a 100X 1.4 NA objective lens, sCMOS camera (Hamamatsu Orca-Flash 4.0 and an iChrome Multi-Laser Engine (Toptica Photonics). The target area was photobleached by continuous excitation at 561 nm prior to acquisition. 5000 frames at 20 ms intervals were acquired on the bleached field of view with continuous activation at 405 nm.

### Single-molecule tracking and data analysis

Bright field images were used to segment cells prior to tracking. Fluorescent spots were detected by a Laplacian of Gaussian (LoG) method in TrackMate (116) and subsequently filtered based on intensity. Spot radii were estimated from the standard deviation of a Gaussian fit. Spots were then linked into tracks using a simple nearest neighbour search and a gap parameter to account for blinking. Tracks shorter than 10 frames were discarded.

Maximum intensity projections were made across 1000 frames, generating 5 projections per movie. Putative clusters were identified in each projection using the LoG method in TrackMate. Individual tracks were assigned to putative clusters if their trajectories overlapped with the cluster area. A threshold of three tracks per cluster was set to remove putative clusters that were defined by a single track. Each molecule was then classified as “In” if its track was localized inside a cluster for its entire duration or “In/Out” if its track was inside a cluster for 50-99% of its duration. All other tracks were classified as “Out”.

For each track, we calculated the time-averaged mean square displacement:

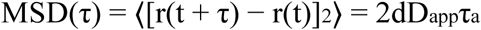

where r(t) is the position at time t, d is the dimension of the system, D_app_ is the apparent diffusion coefficient and a is the anomalous scaling exponent. We extracted D_app_ assuming that a = 1.

## Acknowledgements

We thank Paul Francois, Jie Xiao, Rick Gourse, Jared Schrader and Lisa Racki for thoughtful discussions and members of the Weber and Reyes labs for critical reading of the manuscript. Strain SS6282 was a generous gift from Steven Sandler (UMass-Amherst). The ASKA library was purchased from the National BioResource Project at the National Institute of Genetics (Japan). Deletion strains from the Keio collection were obtained from the E. coli Genetic Stock Center, which is supported by the National Science Foundation (DBI-0742708). This work was supported by the Natural Sciences and Engineering Research Council of Canada (RGPIN-2017-04435 to SCW). A-ML was supported by the Fonds de recherche du Québec - Nature et technologies (B3X-209443).

## Supplementary Figure Legends

**Figure S1.**
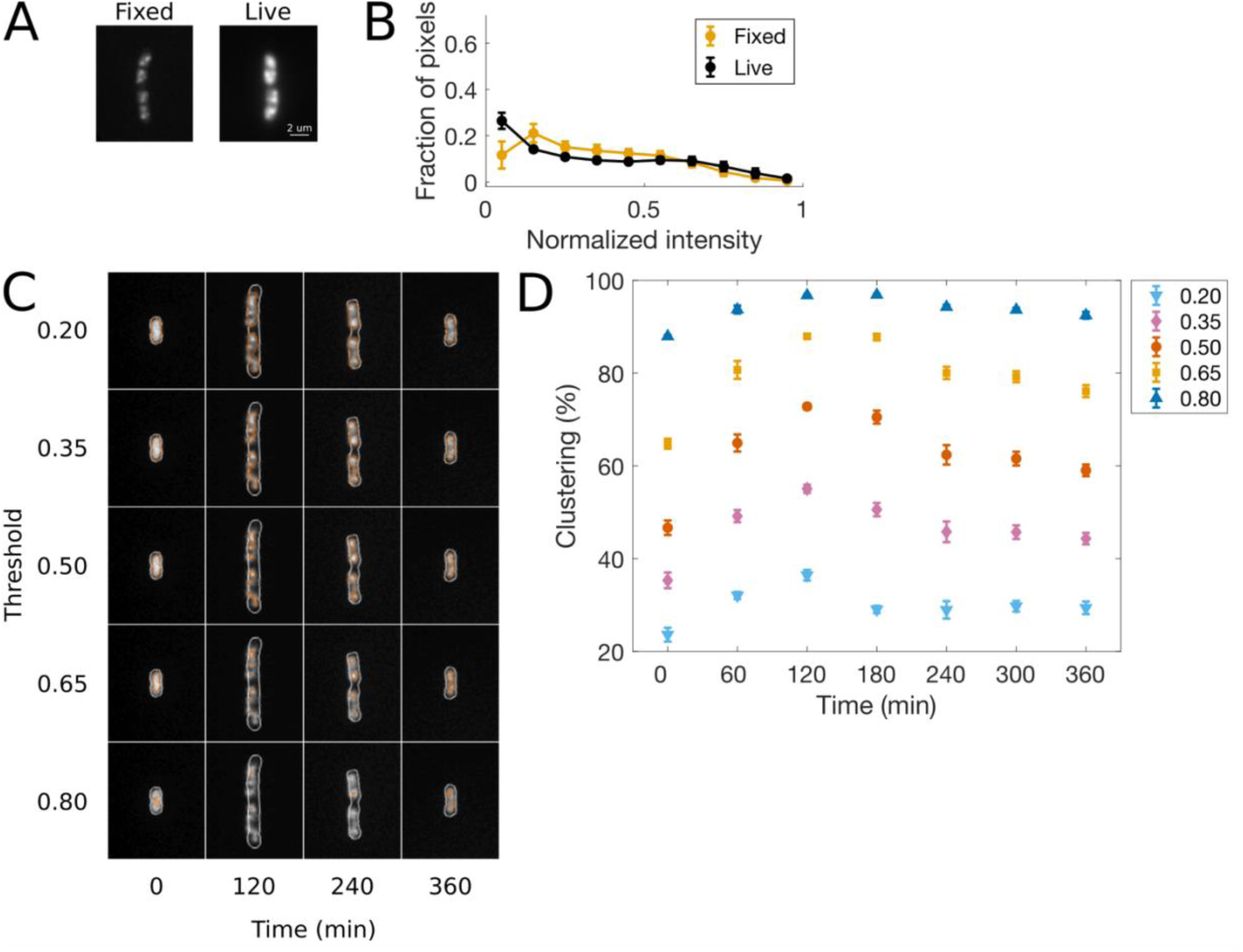
A) Fluorescence images of a fixed cell expressing RpoC-mCherry and a live cell expressing RpoC-mMaple3, both grown in EZ at 37°C for 120 min. B) Normalized pixel intensity histograms for fixed and live cells. C) Fluorescence images of fixed cells grown in LB with cell outlines (white) and segmented RNAP (vermillion) overlaid. The normalized intensity threshold used for segmentation increases from top to bottom. A normalized intensity threshold of 0.50 was chosen because this value best distinguishes between clusters and bulk nucleoid at 120 min. D) Quantification of clustering (percent of pixels below threshold) over time for different threshold values.

**Figure S2.**
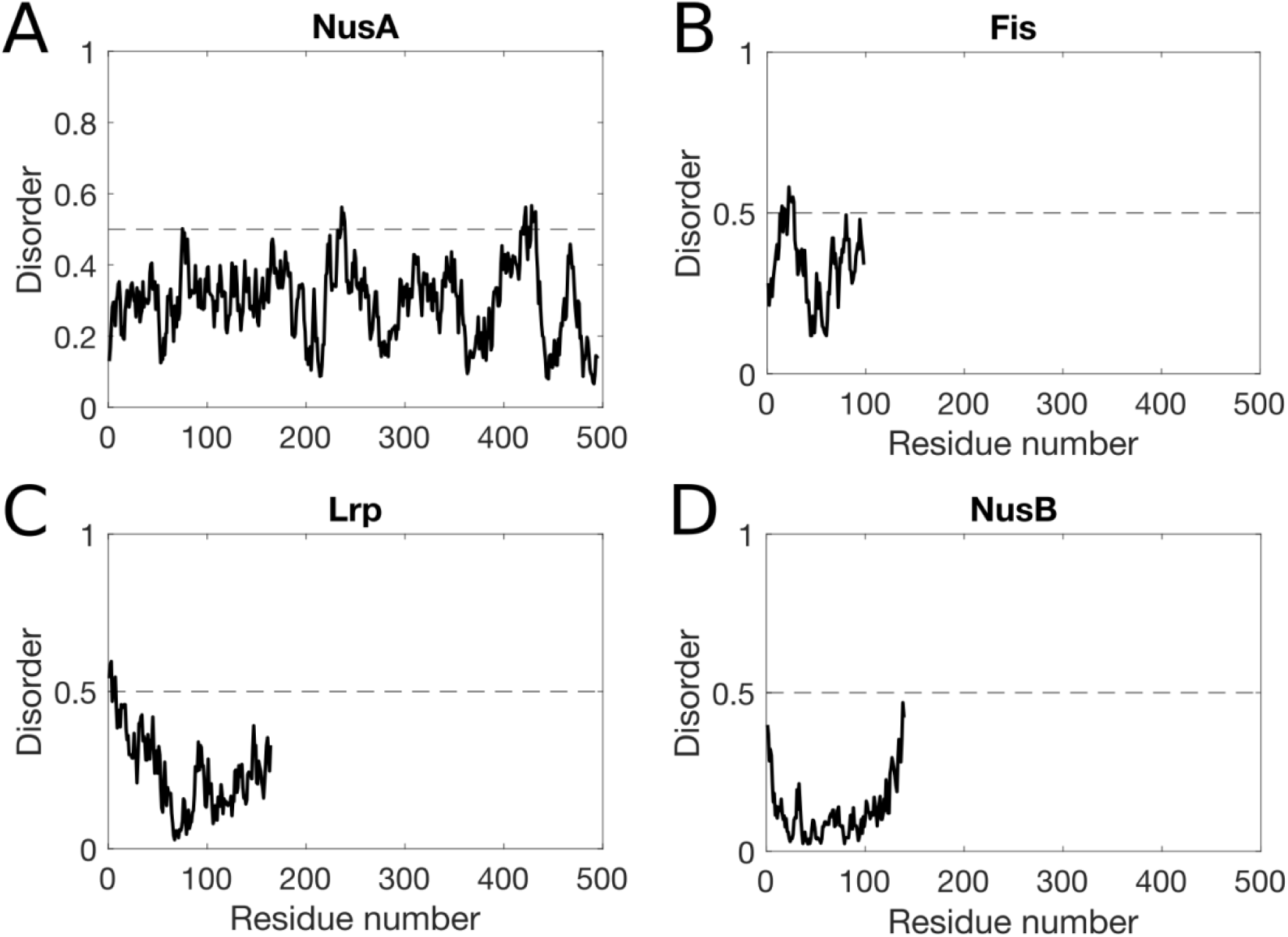
Predicted disorder (by IUPred) for A) NusA, B) Fis, C) Lrp and D) NusB.

**Figure S3.**
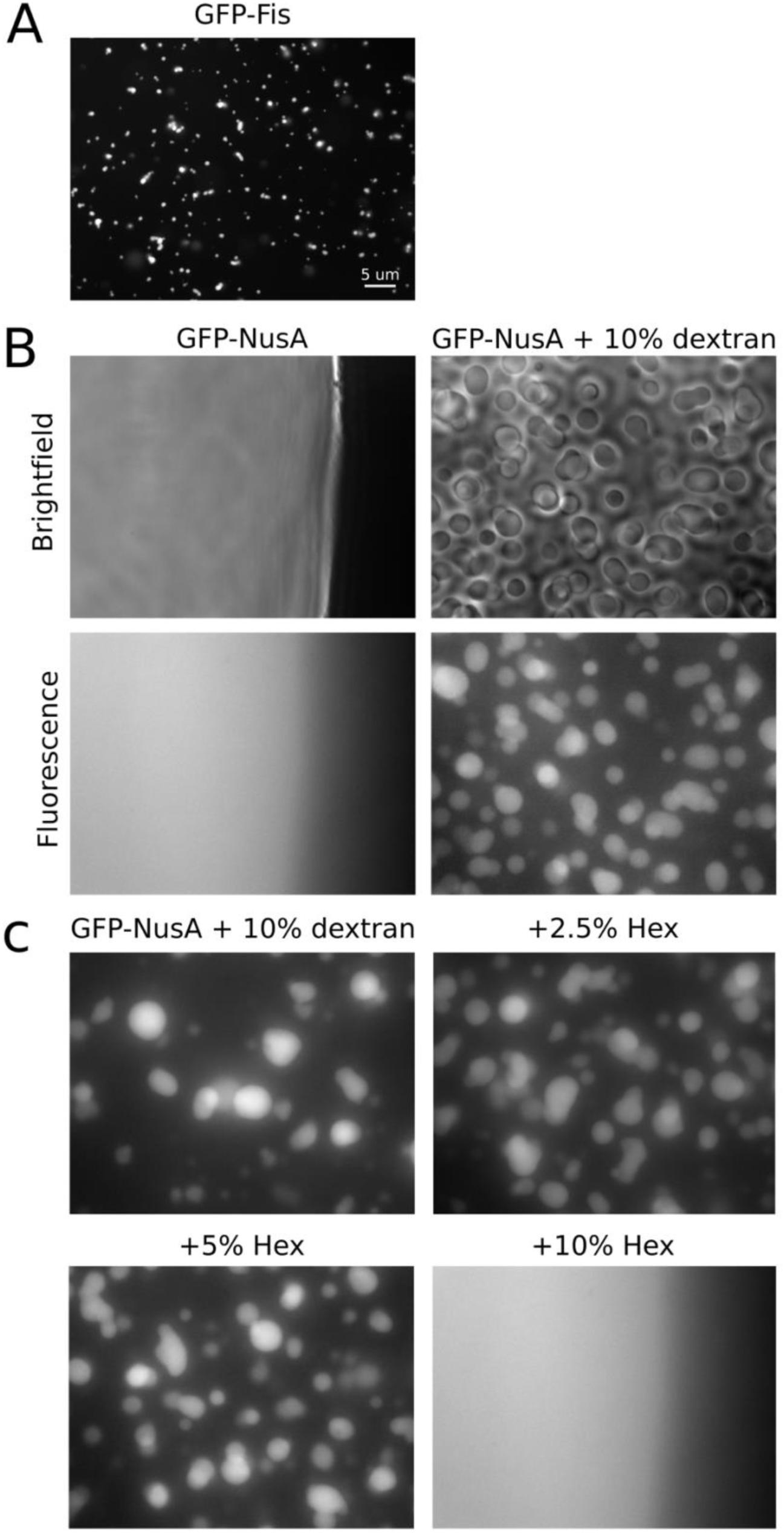
A) GFP-Fis (70 μM) condenses into small droplets at 500 mM NaCl, in the absence of dextran. B) GFP-NusA (14.5 μM) is soluble at 125 mM NaCl in the absence of dextran, but condenses into droplets when 10% dextran is added. C) GFP-NusA droplets (14.5 μM protein, 125 mM NaCl) dissolve in 10% 1,6-hexanediol.

**Figure S4.**
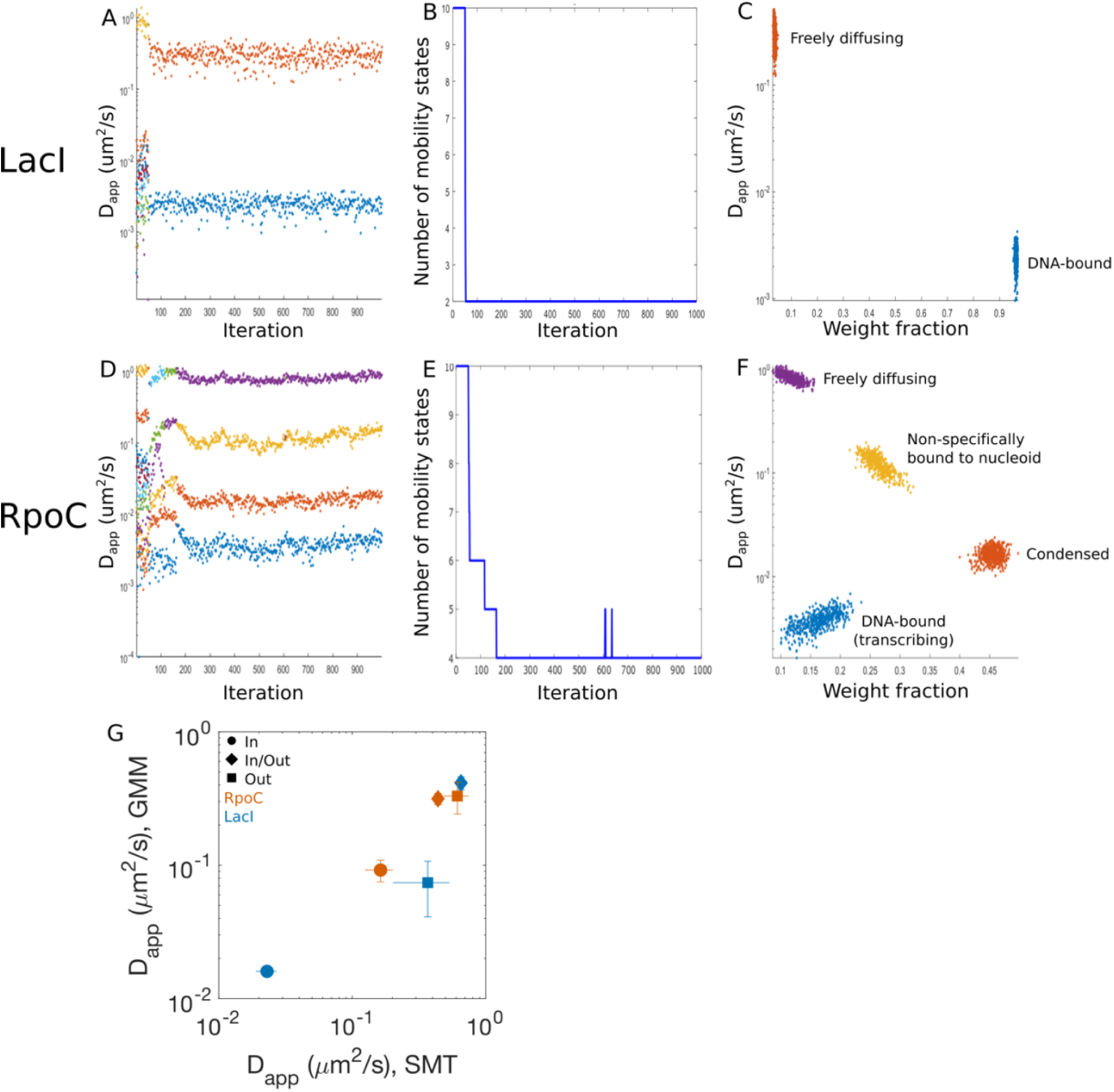
A-C) Results from SMAUG for LacI single-molecule tracking data (in EZ). D-F) Results from SMAUG for RpoC single-molecule tracking data (in EZ). G) Comparison between results from our classification scheme (SMT, described in main text) and a Gaussian mixture model (GMM).

**Figure S5.**
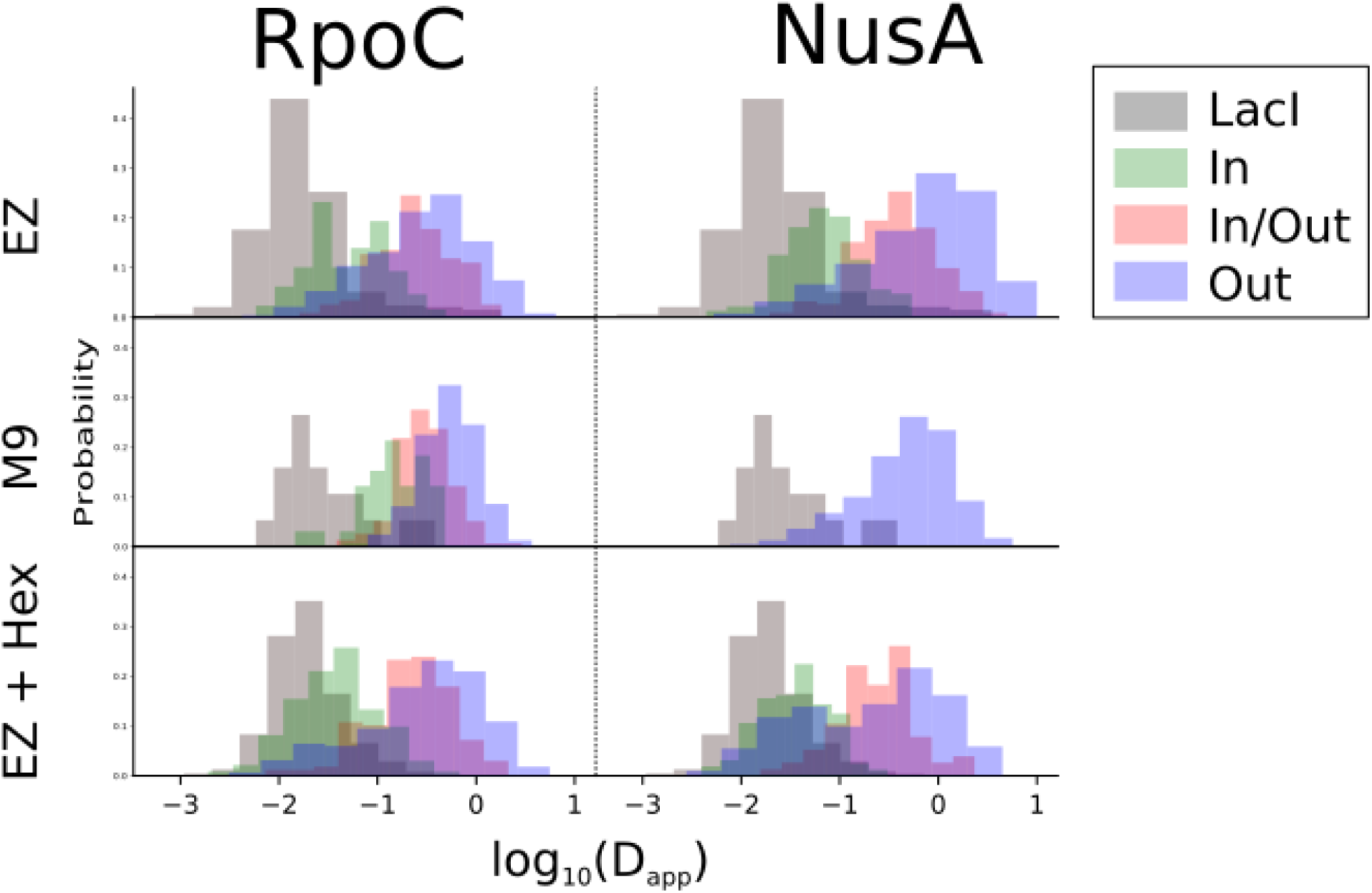
Distribution of D_app_ for LacI, plus each class of RpoC and NusA tracks, in cells grown in EZ, M9 and EZ + 5% 1,6-hexanediol (EZ + Hex).

